# *Hnf1b* renal expression directed by a distal enhancer responsive to Pax8

**DOI:** 10.1101/2022.06.14.496053

**Authors:** L. Goea, I. Buisson, V. Bello, A. Eschstruth, M. Paces-Fessy, R. Le Bouffant, A. Chesneau, S. Cereghini, J.F. Riou, M. Umbhauer

## Abstract

*Xenopus* provides a simple and efficient model system to study nephrogenesis and explore the mechanisms causing renal developmental defects in human. *Hnf1b* (hepatocyte nuclear factor 1 homeobox b), a gene whose mutations are the most commonly identified genetic cause of developmental kidney disease, is required for the acquisition of a proximo-intermediate nephron segment in *Xenopus* as well as in mouse. Genetic networks involved in *Hnf1b* expression during kidney development remain poorly understood. We decided to explore the transcriptional regulation of *Hnf1b* in the developing *Xenopus* pronephros and mammalian renal cells. Using phylogenetic footprinting, we identified an evolutionary conserved sequence (CNS1) located several kilobases (kb) upstream the *Hnf1b* transcription start and harboring epigenomic marks characteristics of a distal enhancer in embryonic and adult renal cells in mammals. By means of functional expression assays in *Xenopus* and mammalian renal cell lines we showed that CNS1 displays enhancer activity in renal tissue. Using CRISPR/cas9 editing in *Xenopus tropicalis*, we demonstrated the *in vivo* functional relevance of CNS1 in driving *hnf1b* expression in the pronephros. We further showed the importance of Pax8-CNS1 interaction for CNS1 enhancer activity allowing us to conclude that *Hnf1b* is a direct target of Pax8. Our work identified for the first time a *Hnf1b* renal specific enhancer and may open important perspectives into the diagnosis for congenital kidney anomalies in human, as well as modeling *HNF1B*-related diseases.

## Introduction

The hepatocyte nuclear factor 1 homeobox B (Hnf1b) is a transcription factor (TF) involved in the development and homeostasis of many organs in vertebrates, including liver, pancreas, hindbrain, genital tract and kidney. In the mammalian embryonic kidney, *Hnf1b* is expressed in tissues leading both to the collecting system and the nephrons. Gene targeting studies have shown that Hnf1b plays a crucial role at several stages of kidney development, including ureteric bud branching and early nephrogenesis as well as nephron patterning and epithelial tubular organization of the collecting ducts ^1–5^. The pivotal role of Hnf1b in renal tubule fate is underscored by the finding that combined expression of the transcription factors Emx2, Hnf1b, Hnf4a and Pax8 is sufficient to reprogram fibroblasts into renal tubular epithelial cells ^6^. Altogether, these studies led to the identification of many Hnf1b target genes and uncovered Hnf1b coordinated transcriptional circuits involved in nephrogenesis and epithelial differentiation ^1–4,7–13^. By contrast, regulation of *Hnf1b* expression in the different embryonic renal compartments remains essentially unknown. Retinoic acid signaling has been shown to directly regulate *Hnf1b* expression in the mouse developing neural tube through a conserved enhancer element but this enhancer is not active in the developing mouse kidney ^14^ and S. Cereghini, unpublished).

Heterozygous mutations in *HNF1B* are one of the most commonly identified genetic cause of developmental kidney disease in human. They are associated with several disorders such as the Renal Cysts and Diabetes Syndrome (RCAD, OMIM #137920), a multi-organ disease, characterized by kidney abnormalities and early diabetes ^15^. Molecular genetic defects comprise whole-gene deletions and intragenic mutations, including missense, frameshift, nonsense, and small insertions or deletions ^16,17^. Mutations are inherited in an autosomal dominant pattern, although up to 50 % of mutations occur de novo ^18,19^. The phenotype of HNF1B mutant carriers is highly variable within families ^19,20^. Consistent with these observations, heterozygous mutants in a novel mouse model carrying a splice donor site mutation identified in humans, exhibited a range of renal phenotypes similar to those described in human and, with some variability, decreased HNF1B protein levels from the normal allele ^11^. Thus, any mechanism that modulates HNF1B expression levels may therefore impact on the variable pathogenic phenotypes in *HNF1B* heterozygous mutant carriers. Identification of the mechanisms underlying *HNF1B* expression is therefore of great importance for the understanding of renal anomalies associated with *HNF1B* mutations. It will potentially uncover novel mutations affecting HNF1B protein levels because they are associated with *HNF1B* regulatory sequences or TFs interacting with these sequences.

*Xenopus* provides a simple and efficient model system to study nephrogenesis and explore the mechanisms that cause renal developmental defects in human ^21,22^.The *Xenopus* tadpole contains only one functional large nephron, the pronephros, displaying a structural and functional organization very similar to the mammalian nephron of the metanephros ^23,24^. Many developmental genes that govern pronephros formation have similar functions during mammalian kidney development. Pax8 has been shown to be a major actor of renal specification and epithelialization in *Xenopus*; its depletion leads to a complete absence of pronephric tubule ^25–27^;. *Pax2* is expressed in the developing pronephros at later stages than *pax8*, once renal cells have been specified, and is required for the expression of several terminal differentiation markers of the pronephric tubule ^25^. The expression of *hnf1b* begins at the late neurula stage in the kidney field and is maintained throughout the entire pronephros, with the highest levels in the proximal region of the pronephric tubule anlage at tailbud stages. Overexpression of an Hnf1b -dominant negative construct in *Xenopus* embryos leads to a strong reduction of the proximal and intermediate pronephric segments and downregulation of Notch ligand *dll1* (*Delta-like1*) and Iroquois *irx1/2/3* gene expression ^3^. These observations are in agreement with the consequences of specific inactivation of *Hnf1b* in mouse nephron progenitors as well as in *Hnf1b*-deficient zebrafish embryos, pointing *hnf1b* as a major regulator of nephron tubular segmentation in vertebrates ^3,10,28^.

Some lines of evidence in *Xenopus* and zebrafish indicate that pax8 and/or pax2 are regulators of *hnf1b* expression. Pax2a/8-deficient zebrafish embryos fail to activate expression of *hnf1ba* in the intermediate mesoderm. In *Xenopus*, pax8 loss of function leads to inhibition of *hnf1b* expression in the kidney field as soon as the late neurula stage while expression of *lhx1* and *Osr2* is not affected. In addition, *hnf1b* is significantly up-regulated in blastula animal caps in response to pax8 ectopic expression even in absence of protein synthesis, suggesting that *hnf1b* might be a direct target of pax8 ^25^. Together, these results suggest a model where pax2 and/or pax8 act upstream of hnf1b, which in turn results in the initiation of segment-specific gene expression programs ^28^.

In the present study, we have explored the transcriptional regulation of *Hnf1b* in the developing *Xenopus* pronephros and mammalian renal cells. Using phylogenetic footprinting, we searched for evolutionary conserved noncoding sequences as potential candidates of *Hnf1b* enhancers. We identified a 306-pb sequence (CNS1) located several kb upstream the *Hnf1b* transcription start site. CNS1 sequence is conserved in *Xenopus*, mouse and human and bears epigenomic marks characteristics of a distal enhancer in embryonic and adult renal cells in mammals. Enhancer activity of CNS1 was established by means of reporter expression assays in *Xenopus* embryos and mammalian renal cell lines. We demonstrated that CNS1 is able to drive reporter gene expression in MDCK and IMCD3 cell lines as well as in the pronephros in *Xenopus* embryos. *In vivo* functional relevance of CNS1 in endogenous *hnf1b* pronephric expression was proven using CRISPR/cas9 in *Xenopus*. Inspection of conserved TF binding sites in CNS1 revealed the presence of a Pax8 binding site. We further demonstrated the importance of Pax8-interaction for CNS1 enhancer activity allowing us to conclude that *Hnf1b* is a direct target of Pax8.

## Results

### Identification of a conserved non-coding sequence as candidate for *hnf1b* enhancer regulated by Pax8

In order to identify regulatory sequences that activate *Hnf1b* gene expression in the developing kidney, we searched for evolutionary conserved non-coding sequences (CNSs) as candidates for enhancers. Using *in silico* methods that allow detailed inspection of sequence conservation profiles across several genomes ^29^, we compared the genomic sequence of a 157 kb segment encompassing the mouse (*Mus musculus*) *Hnf1b* gene with the orthologous intervals in human (*Homo sapiens*), rat (*Rattus norvegicus*), opossum (*Monodelphis domestica*), chicken (*Gallus gallus*), frog (*Xenopus tropicalis*) and zebrafish (*Danio rerio*). This analysis identified one CNS (called CNS1) conserved in all species, except zebrafish, located 33.4 kb upstream the *Hnf1b* transcription start site in mouse and a second one (CNS2) located in the 4^th^ intron of *Hnf1b* conserved in all species examined (Fig. 1a, b). A survey of the literature revealed that CNS2 is identical to a previously characterized *Hnf1b* enhancer whose activity is restricted to *Hnf1b* neural expression in transgenic mice ^14^. We therefore focused our analysis on the 306-bp genomic region CNS1.

**Figure 1.**
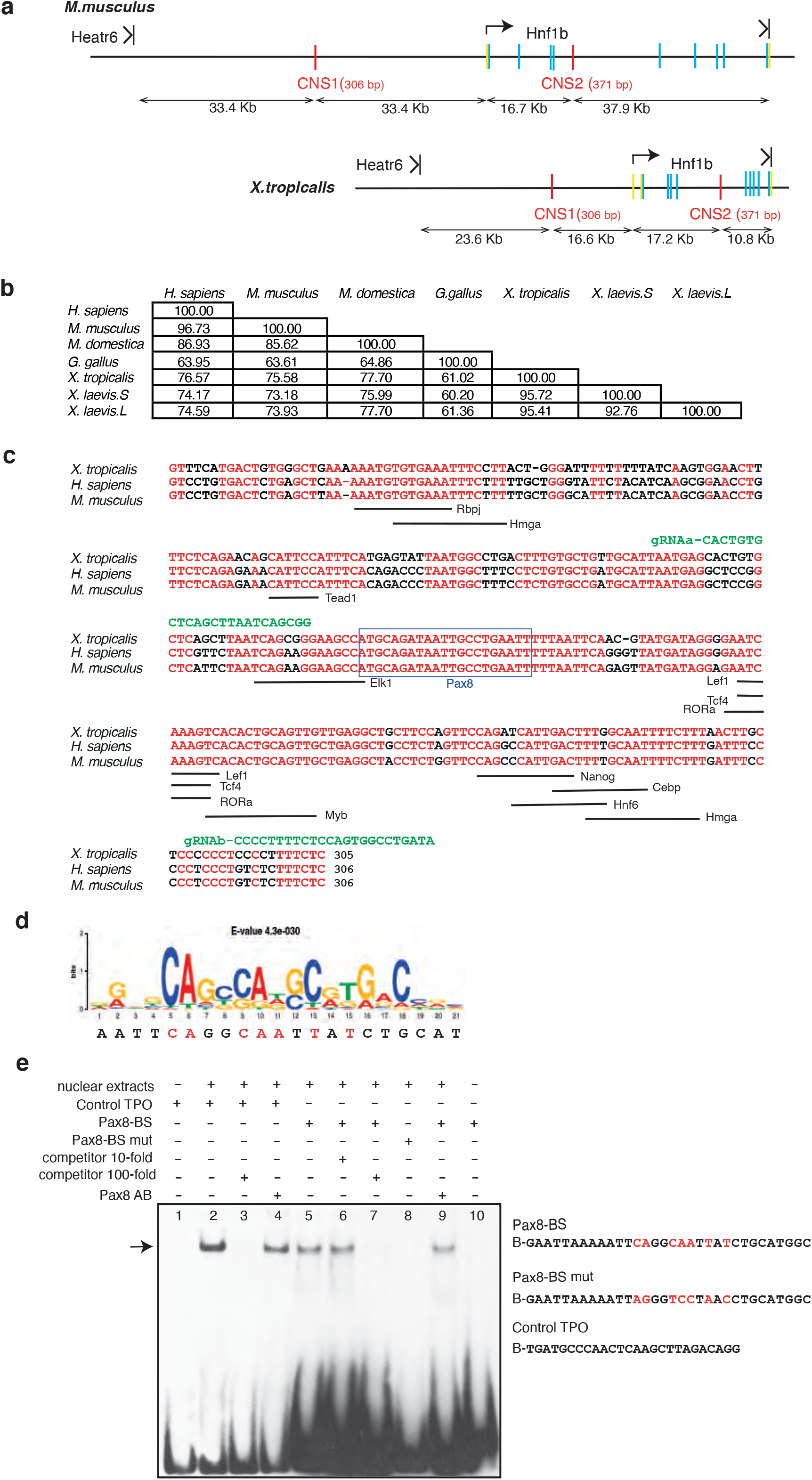
Identification of a conserved non-coding sequence as candidate for *Hnf1b* enhancer regulated by Pax8. **(a)** Localization of two conserved non-coding sequence (CNS1 and CNS2, red bars) in *M. musculus* and *X. tropicalis*. CNS1 is localized in the intergenic region between *Hnf1b* and the upstream gene *Heatr6*. CNS2 is localized in the 4^th^ intron of *Hnf1b*. Arrows indicate the transcription initiation of *Hnf1b*. Protein coding exons are represented by blue bars, exonic sequences corresponding to 5’and 3’ UTRs are in yellow. **(b)** Nucleotide sequence comparison of CNS1 in different species. The table indicates the % of conserved nucleotides. **(c)** Nucleotide sequence alignment of CNS1 in *X. tropicalis, H. sapiens* and *M. musculus*. Conserved nucleotides are in red. Underlined are conserved putative binding sites of TFs known to be expressed in the developing kidney according to ^74^ (https://sckidney.flatironinstitute.org), GUDMAP (https://www.gudmap.org) or Xenbase (https://www.xenbase.org). The blue box highlights the conserved Pax8 putative binding site. Sequences targeted by the gRNAa and gRNAb used for CRISPR editing in figure 6 are shown in green. **(d)** Pax8 binding site sequence identified in CNS1 and Pax8 binding consensus sequence as defined by shown as sequence logo. For each position, the size of the characters represents the relative frequency of the corresponding position. The lower sequence is the Pax8 binding sequence in CNS1. Nucleotides mutated in the Pax8-BS mut oligonucleotide used for Electrophoretic Mobility Shift Assay (EMSA), and in plasmids CNS1 mut-Luc and CNS1 mut-eGFP are labelled in red. **(e)** EMSA. Biotin-labelled oligonucleotide probes were incubated with nuclear extracts from MDCK as indicated: Pax8-BS, Pax8-BS mut and Control TPO (Pax8 binding site identified in the thyroperoxidase promoter, used as a positive control). An arrow marks the Pax8 specific complexes. Specificity of the Pax8 complex was demonstrated by competition with 10- and 100-fold molar excesses of unlabelled Pax8-BS oligonucleotide (compare lanes 5 and 7) as well as incubation with 1μg of Pax8 monoclonal antibodies (Pax8 AB) leading to a decrease in the Pax8 complex signal (compare lanes 5 with 9). The figure is representative of 3 independent experiments. It presents the autoradiography of the whole blot membrane.

Active regulatory sequences show epigenomic properties such as characteristic patterns of histone modification and accessible chromatin. CNS1 was queried against public databases for Deoxyribonuclease I (DNase I) hypersensitive site sequencing (DNase-seq), Assay for Transposase-Accessible Chromatin sequencing (ATAC-seq) and chromatin-immunoprecipitation sequencing (ChIP-seq) for histones marks or the transcriptional cofactor p300 from human, mouse (ENCODE project) and *Xenopus* ^30^ embryonic and adult tissues or cell lines. We found that CNS1 correlates with a DNase I hypersensitive site in several human primary cells from adult kidney tubule known to express HNF1b at a high level but not in Caco-2 or A549 cell lines derived from colon carcinomas and lung adenocarcinomas respectively, which also express HNF1b although at a lower level (https://www.proteinatlas.org) (Fig. 2a; Fig. S1a). No DNase I hypersensitive signal was detected at the CNS1 location in spinal cord or neural progenitors (Fig. 2a; Fig. S1a). In the mouse embryo, *Hnf1b* is expressed during kidney development but also in the developing liver, pancreas and lung ^31–33^. We looked for CNS1 epigenetic status in these embryonic tissues. We observed that CNS1 showed characteristics of an active enhancer in E15.5 embryonic kidney: it overlaps with an ATAC-seq signal which is bordered by H3K4me1, H3K4me2 and H3K27ac signals. Interestingly, no such signal was observed in lung or liver tissue (Fig. 2b, c; Fig. S1b). In *Xenopus*, we also found CNS1 associated with active enhancer marks (p300, H3K4me1) in ChIP-seq data from whole neurula and tailbud stages embryos (stages 16 and 30, respectively) when *hnf1b* is expressed in the developing pronephros (Fig. 2b, d; Fig. S1c).

**Figure 2.**
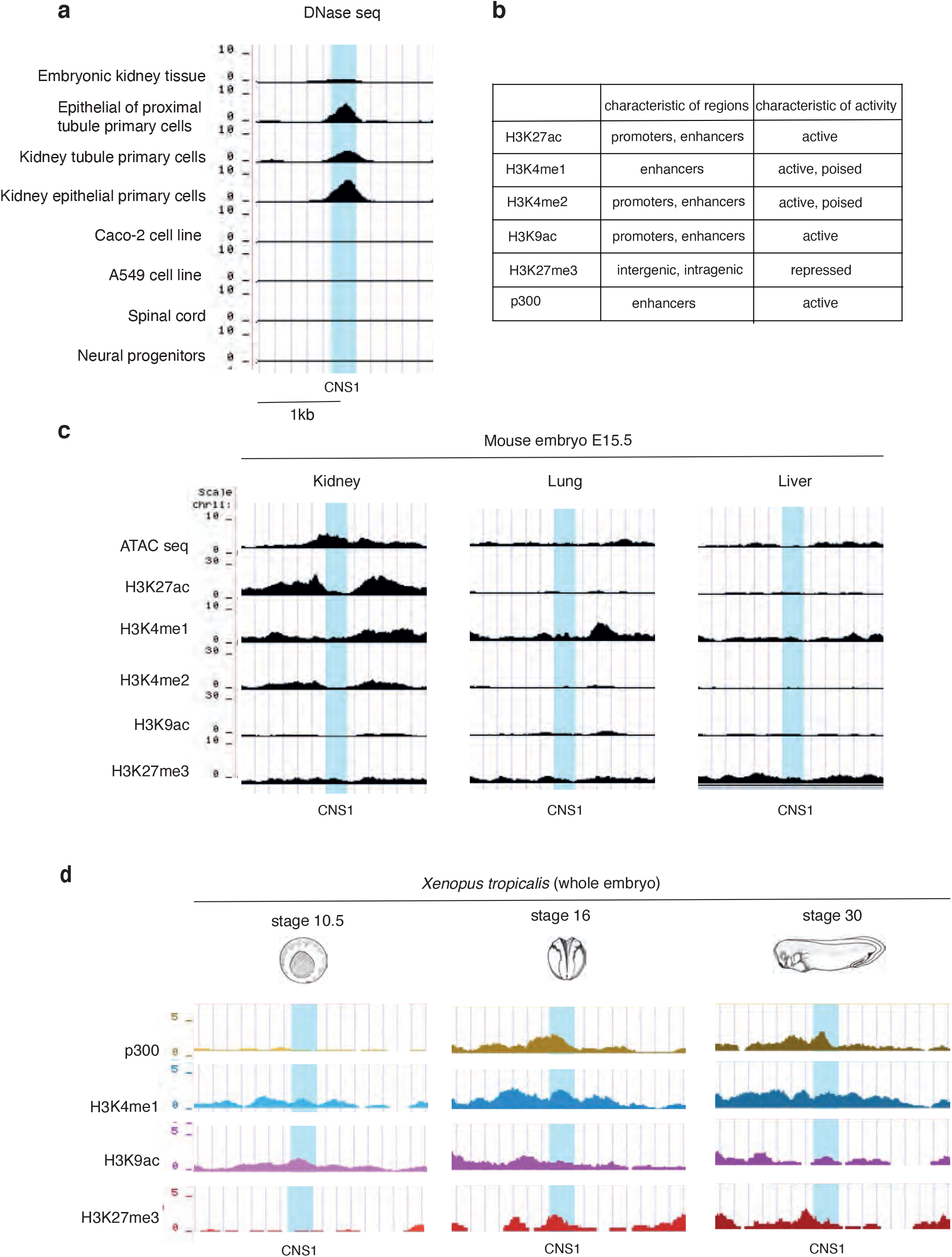
Chromatin signature of CNS1 region in *Homo sapiens, Mus musculus* and *Xenopus tropicalis*. (**a**) Snapshots of representative DNase-seq tracks from ENCODE database in human (https://www.encodeproject.org). The overall profiles spanning a larger region including CNS2 and the description of the biological materials are shown in Fig. S1a. Open chromatin state is clearly detected at CNS1 level in embryonic kidney tissue and renal cells. In contrast, CNS1 is not accessible to DNase in non-renal cells expressing HNF1B (Caco-2 and A549 cells) as well as in spinal cord or neural progenitors where HNF1B is not expressed. (**b**) Characteristics of enrichment patterns for histone modifications and p300. (**c**) Snapshots of representative ATAC-seq and histone modifications tracks as indicated from ENCODE in kidney, lung and liver of E15.5 mouse embryo. The overall profiles spanning a larger region including CNS2 is shown in Fig. S2b. Although Hnf1b is expressed in developing kidney, liver and lung, CNS1 shows characteristics of an active enhancer only in the developing kidney. **(d)** Snapshots of representative p300 and histone modifications tracks as indicated from the epigenome reference maps in ^30^ in *Xenopus tropicalis* embryos at stages 10.5, 16 and 30. The overall profiles spanning a larger region including CNS2 is shown in Fig. S1c. CNS1 is associated with active enhancer marks (p300, H3K4me1) in embryos at neurula and tailbud stages (stage 16 and 30, respectively) when *hnf1b* is expressed in the developing pronephros but not at early gastrula (stage 10.5). In a, c and d, the blue box indicates the 306-bp CNS1 location.

Altogether, these data suggest that CNS1 is an enhancer whose activity is potentially specific to renal tissue. Notably, CNS2 is not localized in a DNase I hypersensitive site in any the analyzed cell lines and does not corelate with H3K27ac enrichment in mouse embryonic kidney (Fig. S1a,b).

We then searched for conserved TF putative binding motifs in CNS1 using R-Vista 2.0 ^34^. We identified 23 putative TF binding sites conserved between frog, mouse and human, including Pax8 whose knockdown in *Xenopus* leads to inhibition of *hnf1b* expression in the kidney field ^25^. Eight additional ones were conserved solely between frog and human (Table S1, S2). The sequence of the Pax8 putative binding site in CNS1 shows high similarity to known Pax8 binding sites ^35^ and is strictly conserved in mouse and human CNS1 (Fig. 1c,d). The actual binding of Pax8 to the identified sequence was tested by electrophoretic mobility shift assays (EMSAs). As a positive control we used a known Pax8-binding site derived from the thyroperoxidase promoter, a target gene of PAX8 ^36^ (Control TPO, Fig. 1e). When incubated with nuclear extracts from Madin-Darby Canine Kidney (MDCK) cells containing abundant levels of Pax8 protein ^37^, biotin-labelled Pax8-BS and control-TPO oligonucleotides formed a similar protein-DNA complex, as judged by their identical electrophoretic mobility (Fig.1e). When excess unlabelled wild-type oligonucleotides were added as competitors, they weakened or abolished completely the shifted bands (Fig. 1e, compare lanes 2 and 3 for control TPO and lanes 5, 6 and 7 for Pax8-BS). No complex was detected with biotinylated oligonucleotides containing mutations known to abolish Pax8 binding ^38^. The presence of Pax8 antibodies in the reaction did not induce a supershift, but decreased the signal intensity of the electrophoretic complexes, suggesting that antibody interaction disrupts Pax8 interaction with both control-TPO and Pax8-BS oligonucleotides (Fig. 1e, compare lanes 2 and 4 for control TPO and lanes 5 and 9 for Pax8-BS), further confirming the specificity of Pax8 binding. Together, these data show that Pax8 is able to specifically bind *in vitro* to the *in silico* identified Pax8 binding site of CNS1.

### CNS1 is specifically activated by ectopic expression of pax8 and pax2 and functions as an enhancer in mammalian renal cells

We subsequently tested the ability of different versions of exogenous pax8 to transactivate CNS1 in HEK293 cells that do not express pax8 protein (www.proteinatlas.org/ENSG00000125618-PAX8/cell) using luciferase assays. The CNS1 sequence was cloned into the luciferase pGL4.23[*luc2*/minP] vector and the resulting construct CNS1-Luc was used to transfect HEK293 cells together with a control vector carrying *Renilla* luciferase gene used to normalize for transfection efficiency. We analysed the effect of expressing wild-type pax8, pax8ΔO, a mutated version of pax8 the octapeptide motif mediating interaction with Groucho/Grg4 co-repressors responsible for transcriptional repressor activity ^39,40^, and pax8VP16, a chimeric protein corresponding to pax8 fused to the potent transcriptional activation domain of the herpes simplex virus VP16 and conferring strong transcriptional activity to pax8 ^25^. HEK293 cells were transfected with CNS1-Luc vector alone or associated with one of the pax8 expression vectors (pax8, pax8VP16, pax8ΔO). Expression of either pax8, pax8VP16 or pax8ΔO led all to an increase in relative luciferase activity. As expected, this increase was much higher for pax8△O and pax8VP16 (~40-fold) than for wild-type pax8 (~5-fold), and further suggested that Groucho/Grg4 repression is also at work in HEK293 cells (Fig. 3a-b). Mutation of the Pax8 binding site in CNS1 almost abolished this activation in every instance (Fig. 3a-b). Our results therefore show that CNS1 transactivation in HEK293 cells is dependent on the expression of exogenous Pax8, and the integrity of a sequence that binds Pax8 in *vitro*, strongly suggesting that this process results from the interaction of exogenous Pax8 with this sequence.

**Figure 3.**
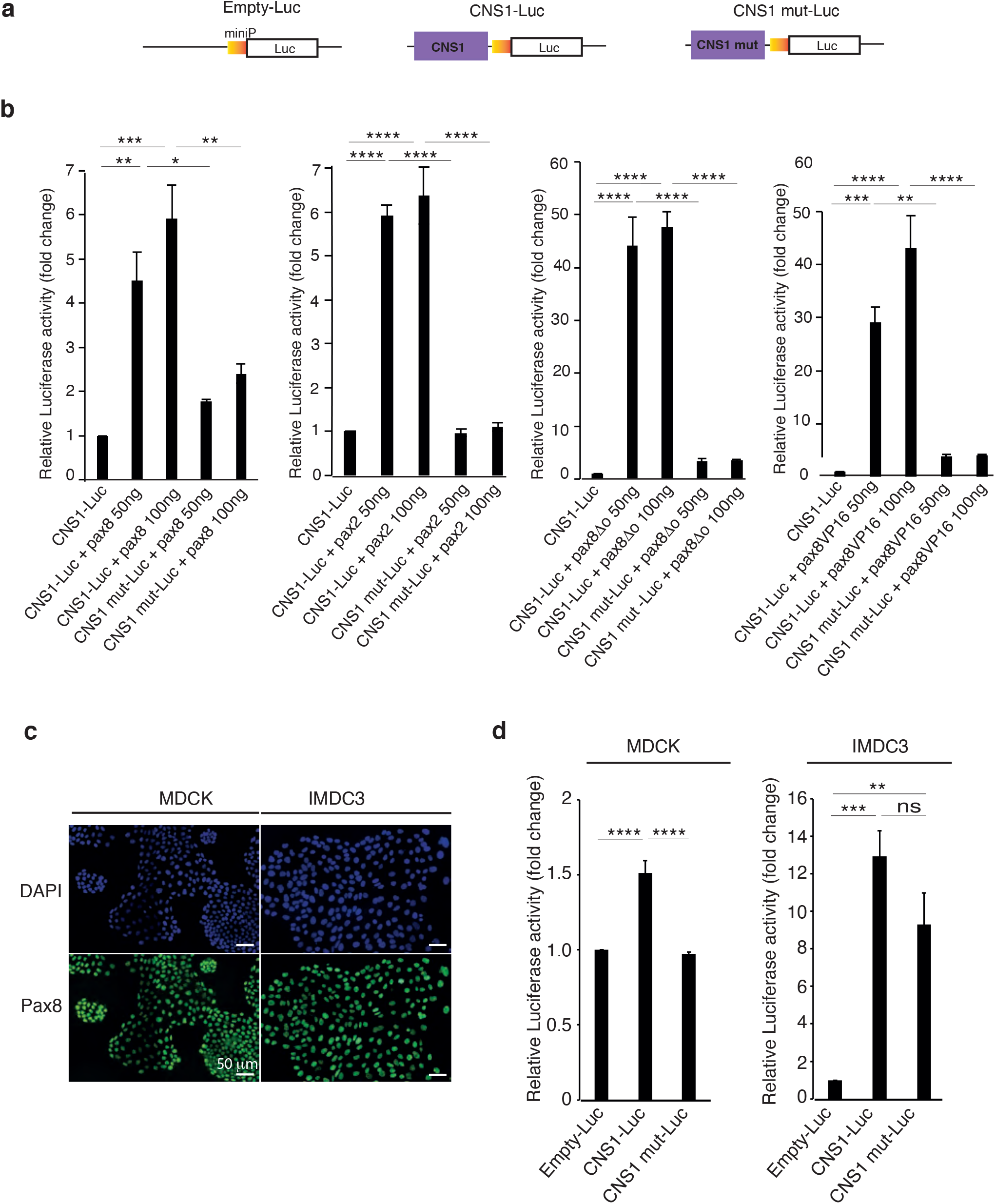
CNS1 is specifically activated by ectopic expression of pax8 and pax2 and functions as an enhancer in mammalian renal cells. **(a)** Reporter constructs used in luciferase assay experiments. The backbone plasmid pGL4.23[*luc2miniP*] was used either unmodified (Empty-Luc) or modified by insertion of the wild type CNS1 (CNS1-Luc) or CNS1 mutated in the Pax8 binding site (CNS1 mut-Luc) upstream of the minimal promoter. **(b)** Exogenous pax8 and pax2 are able to activate CNS1 enhancer activation in HEK293 cells. Expression vectors for pax8, pax2, pax8ΔO or pax8VP16 were co-transfected with CNS1-Luc or CNS1mut-Luc, and a control Renilla luciferase vector for normalization. Normalized luciferase activity was determined as the Firefly over Renilla luciferase activity. Fold change is expressed as the ratio of normalized values on the normalized value for CNS1-Luc vector alone. Expression of pax8, pax8ΔO, pax8VP16 and pax2, all result in an increase of reporter activation. In every case, this increase is abolished when the mutant construct, CNS1mut-Luc, is used instead of CNS1-Luc. Combined results from three (pax8, pax8ΔO and pax8VP16) or four (pax2) independent experiments. *\*p<0.05, **p<0.01, ***p<0.001, ****p<0.0001*. Statistical significance was determined using One-Way ANOVA followed by Tuckey’s multiple comparison test **(c)** Pax8 immunofluorescence staining in MDCK and IMCD3 cells and counterstaining of nuclei with DAPI. Pax8 is detected in MDCK and IMCD3 cell nuclei. Scale bar: 50 μm. **(d) (c)** Analysis of CNS1 activity in luciferase reporter assay in MDCK and IMCD3 cell lines. Histograms show the relative luciferase activities in MDCK and IMCD3 cells transfected with Empty-Luc, CNS1-Luc or CNS1 mut-Luc together with a control Renilla luciferase vector for normalization. Fold change is expressed as the ratio of normalized values on the normalized value for Empty-Luc. Combined results from four independent experiments. *\*\*p<0.01, ***p<0.00, ****p<0.0001*. Statistical significance was determined using One-Way ANOVA followed by Tuckey’s multiple comparison test.

The vertebrates *PAX* genes are classified into four sub-groups and *Pax8* belongs to subgroup 2 together with *Pax2* and *Pax5* exhibiting similar recognition sequences. Since *Pax2*, but not *Pax5*, is expressed in the developing kidney ^41^ (GUDMAP https://www.gudmap.org), we evaluated the ability of pax2 to increase transcriptional activity of the CNS1-Luc reporter construct in HEK293 cells. Pax2 expression induced luciferase activity from the CNS1-Luc construct and was unable to activate transcription from CNS1 mut-Luc (Fig. 3a-b). These results indicate that both pax2 and pax8 are able to activate CNS1 in HEK293 cells when ectopically expressed.

We then investigated the functionality of the identified CNS1 in the kidney derived cell lines MDCK and IMCD-3, which both express Pax8 (Fig. 3c) as well as Hnf1b ^37,42^. Luciferase activity of CNS1-Luc transfected cells was significantly increased compared to cells transfected with the empty luciferase vector pGL4.23[*luc2*/minP] (Empty-Luc) in both cell lines (Fig. 3d) although to a greater extend in IMCD3 in comparison to MDCK cells. Mutating Pax8 binding site led to a significant decrease of reporter activity in transfected MDCK cells but not in IMCD3 (Fig. 3d).

Thus, CNS1 enhancer is active in mammalian renal cell lines and its dependency upon the Pax8 binding site varies between cell lines.

### CNS1 functions as a nephric enhancer *in vivo* in *Xenopus* embryos and its activity is partly dependent on the transcription factor pax8

CNS1 enhancer activity was then investigated *in vivo* in *Xenopus laevis* by means of F0 transgenesis ^43^. A plasmid containing CNS1 upstream of a beta-globin basal promoter and eGFP coding sequence was used to generate transgenic embryos through the *I-SceI* meganuclease method ^44^. Reporter expression was examined at different developmental stages by *in situ* hybridization for eGFP mRNA. The control reporter construct without CNS1 (beta-globin-eGFP), did not lead to any significant GFP expression at all stages examined (Fig. 4a, c, d). In contrast, CNS1 drove strong and reproducible expression in the developing pronephros starting at the mid tailbud stage 25 (Fig. 4a-d). Transverse sections showed expression in the developing tubule anlage but not in the splanchnic mesoderm from which the glomus forms (Fig. 4c). GFP reporter expression was therefore similar to the endogenous *hnf1b* expression in the developing pronephros except that endogenous *hnf1b* mRNA is detected earlier, as soon as the late neurula stage 19 in the kidney field (Fig.S2). GFP expression was additionally observed in posterior somites in CNS1-eGFP embryos, although somites do not express endogenous *hnf1b* (Fig. S3). Using the REMI transgenesis method that allows the production of fully transgenic-non-mosaic embryos with high efficiency ^45^, we confirmed that CNS1 was able to drive reporter expression in the pronephros, as shown by GFP fluorescence in the pronephros in 32.6% of the injected embryos (Fig. 4e).

**Figure 4.**
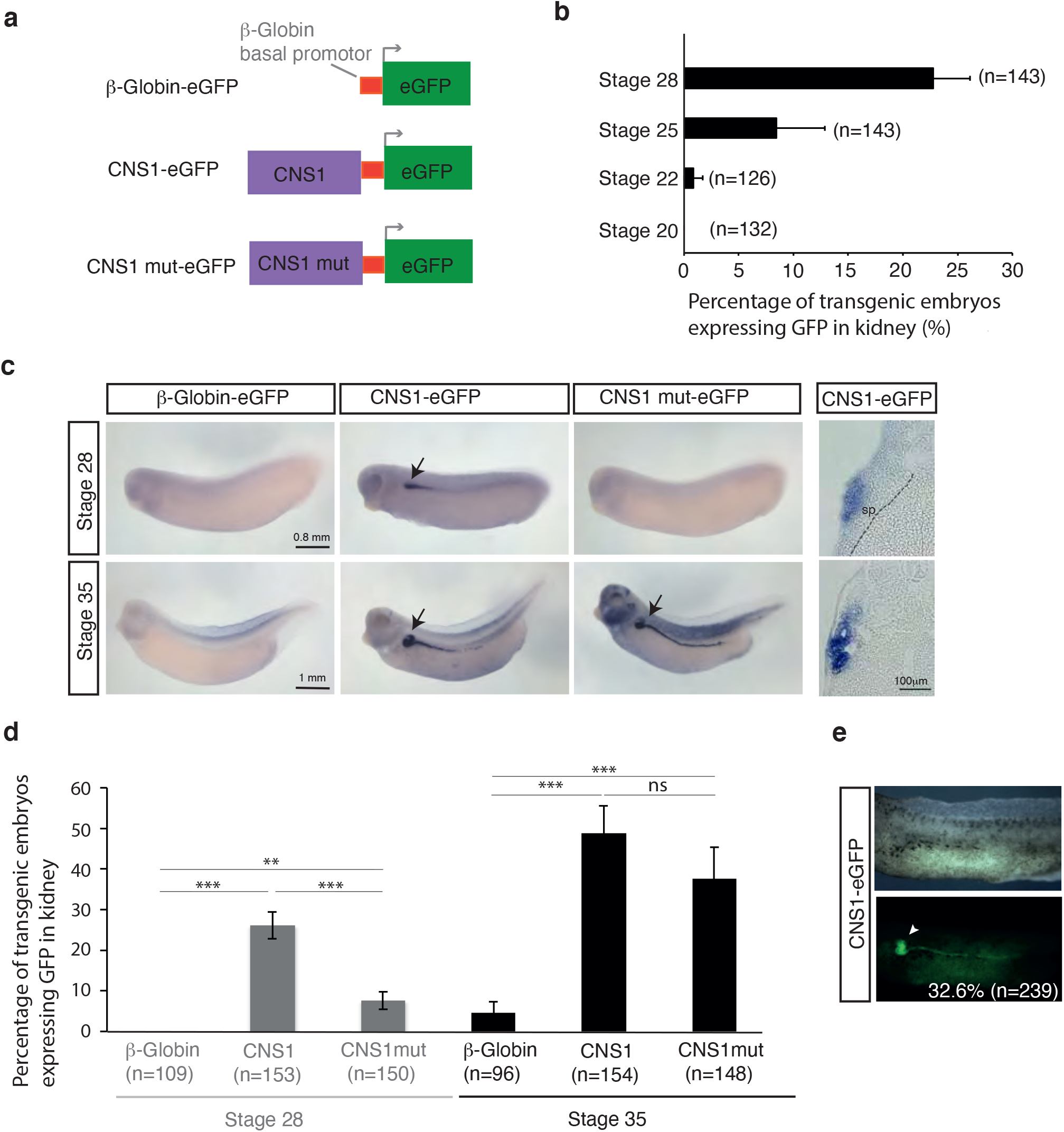
Analysis of CNS1 activity in transgenesis assay in *Xenopus laevis* **(a)** Reporter constructs used in transgenesis. Wild type CNS1 or CNS1 mutated in the Pax8 binding site (CNS1 mut) was cloned upstream of the β-globin basal promoter and the eGFP reporter gene **(b)** Percentage of F0 CNS1-eGFP transgenic embryos generated by the *I-Sce*I meganuclease method showing GFP expression detected by *in situ* hybridization in the developing pronephros at stages 20, 22, 25 and 28. Results corresponding to three independent experiments. **(c)** GFP expression detected by *in situ* hybridization in representative F0 transgenic embryos at stages 28 and 35 generated with reporter constructs as indicated. Transverse cryostat sections (18 μm) of CNS1-eGFP transgenic embryos showing GFP reporter gene expression at the level of the proximal part of the pronephros are shown on the right. At stage 28, the dotted line delineates the mesoderm-endoderm frontier. sp : splanchnic mesoderm **(d)** Percentage of F0 transgenic embryos generated with the indicated reporter constructs expressing GFP in pronephros at stages 28 and 35. Results from three independents experiments. Statistical significance was determined using Fisher’s exact test. *\*p<0.05, **p<0.01, ***p<0.001*. **(e)** Representative stage 40 F0 transgenic tadpole obtained by the REMI method with the CNS1-eGFP construct. Arrowhead indicates the fluorescent pronephros. In b, d and e, n indicates the total number of analysed embryos.

Mutation of the Pax8 binding site greatly diminished the frequency of pronephric GFP expression at stage 28 when compared to the wild-type CNS1-eGFP construct (Fig. 4c, d). However, GFP expression at stage 35 was not affected, although *pax8* is known to be expressed in the pronephric tubule at this stage ^41^ (Fig. 4c, d). These results suggest that additional inputs are present in *Xenopus* embryos at stage 35 that can overcome the mutation of the Pax8 binding site for CNS1 activity.

*Pax2* and *pax8* are both expressed in the developing pronephros at tailbud stage 28, but, in contrast to pax8, pax2 is not required for *hnf1b* expression at tailbud stage ^25^(Fig. 5a). Although unlikely, it is still possible that in stage 28 transgenic embryos pax2 may be involved in CNS1-driven reporter activation instead of pax8. We therefore monitored the effect of pax2 depletion upon reporter gene activity in CNS1-eGFP transgenic embryos. After injecting CNS1-eGFP construct at the 1-cell stage for transgenesis, a pax2 antisense morpholinos (Mo pax2) was injected at the 2-cell stage into the two blastomeres, and reporter expression analysed at stage 28 (Fig.5b). For each experiment, the efficiency of Mo pax2 was confirmed by verifying inhibition of *clcnkb1* expression ^25^ in a separate group of embryos cultured until stage 35 (not shown). Comparison of the percentage of embryos showing GFP mRNAs expression in the developing pronephros of embryos injected with Mo pax2, or with a control morpholino (cMo), did not show any difference (Fig. 5b). These observations show that endogenous pax2 is not required for CNS1 activity in the pronephros at tailbud stage 28. In contrast, when morpholinos targeting pax8 (Mo pax8) were used, the vast majority of injected embryos did not show any GFP signal (Fig. 5b). It is unclear however if this results from the absence of pax8 protein binding to CNS1, or because early development of the pronephric tubule is severely affected by pax8 depletion as previously shown ^25^.

**Figure 5.**
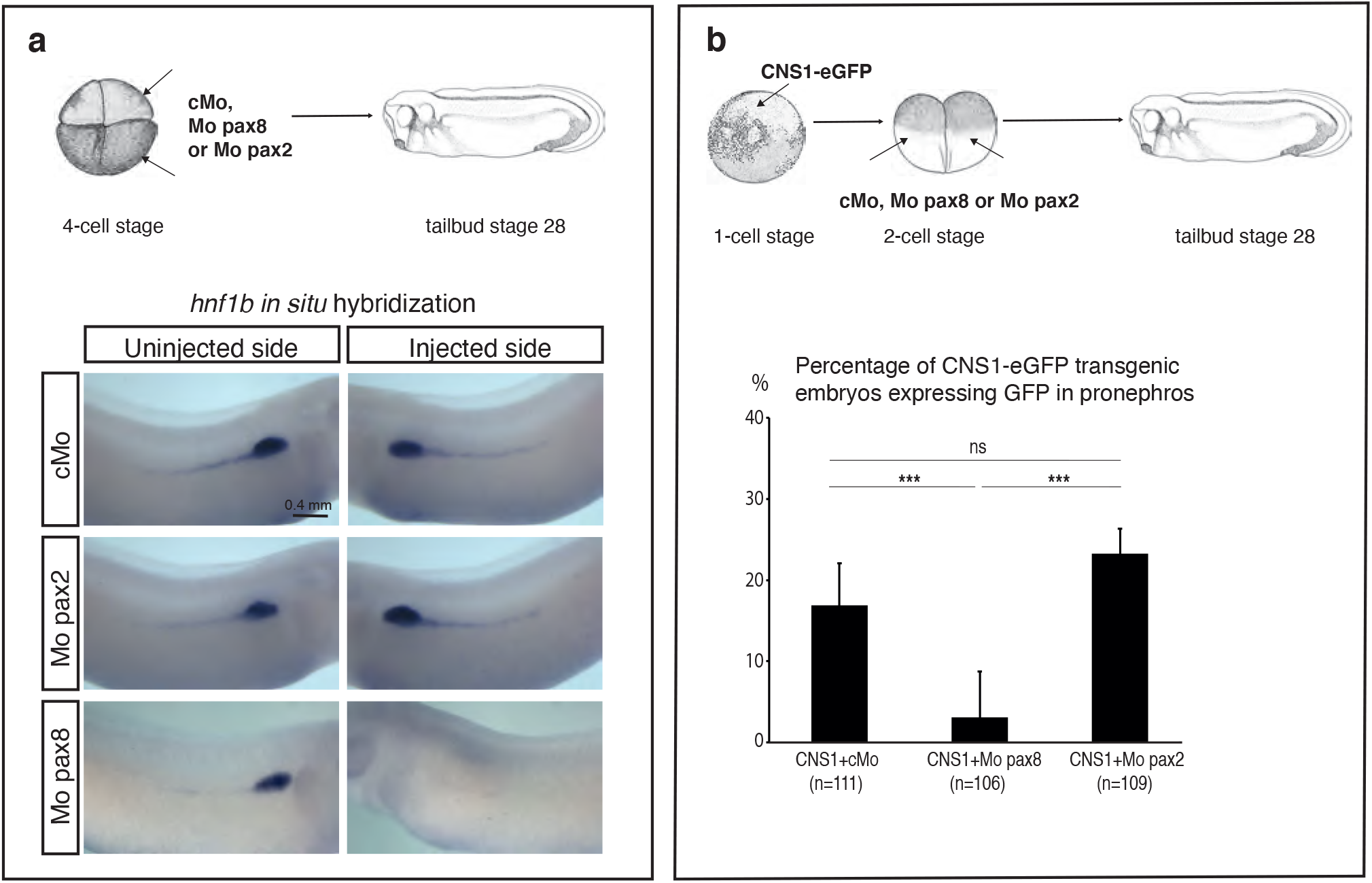
Endogenous pax8, but not pax2, is required for *hnf1b* expression and CNS1 activity in the pronephros at tailbud stage. **(a)** Pax2 depletion does not affect *hnf1b* expression at stage 28. 4-cell stage embryos were injected into the two left blastomeres with either a control Mo (cMo), a mix of two morpholinos, Mo Pax8 and Mo Pax8.B, (named Mo pax8 for simplicity) or Mo Pax2.2 (named Mo pax2). They were further cultured until tailbud stage 28 and processed for *hnf1b* whole-mount *in situ* hybridization. *Hnf1b* pronephric expression is strongly inhibited in 80% (n=24) of pax8 depletion. In contrast, pax2 depletion has no significant effect on *hnf1b* pronephric expression (n=60), as embryos injected with cMo (n=71) (three independent experiments) **(b)** Pax2 depletion does not affect CNS1-driven GFP expression in the pronephros. F0 CNS1-eGFP transgenic embryos were generated by the *I-SceI* meganuclease method at the 1-cell stage. At the 2-cell stage, same mixes of Mo as in (a) were injected into every blastomeres. GFP expression was monitored by whole-mount *in situ* hybridization in embryos cultured until the tailbud stage 28. The histogram indicates the percentage of CNS1-eGFP transgenic embryos showing GFP expression in controls, or upon depletion of pax2 or pax8. In contrast to pax8 depletion, pax2 depletion has no significant effect on CNS1 activity. Statistical significance was determined using Fisher’s exact test. ns: not significant, *\*\*\*p<0.001*. Combined results from three independent experiments. n indicates the total number of analysed embryos.

Altogether, these observations argue for a role of CNS1 in pax8 regulation of *hnf1b* at tailbud stage but also indicates that this enhancer is likely to be regulated by additional not yet identified inputs at later developmental stages.

### CNS1 is required for endogenous *hnf1b* expression in the developing pronephros

In order to evaluate the contribution of CNS1 in its normal genomic environment to the regulation of endogenous *hnf1b* expression, we attempted to mutate CNS1 using the available CRISPR/Cas9 genome editing approach in *Xenopus tropicalis*. We designed two gRNA (gRNAa and b) targeting two sequences encompassing a 180 bp region of CNS1 that includes the pax8 binding sequence (Fig. 1c). TIDE analysis indicated that gRNAa and gRNAb efficiently target CNS1 in F0 embryos (Fig.6a). We co-injected gRNAa and gRNAb with Cas9 protein in *Xenopus tropicalis* embryos with the objective to delete the CNS1 DNA sequence comprised between sgRNAa and sgRNAb targeted sites. Electrophoresis analysis of PCR products amplified with primers surrounding CNS1 revealed a lower band corresponding to a truncated sequence in 3 out of 5 embryos, indicating that the deletion of the targeted sequence occurred with high frequency (Fig. 6b). Embryos were injected with gRNAa and gRNAb alone or in combination, reared to tailbud stage 28 and analyzed for *Hnf1b* pronephric expression by *in situ* hybridization. About 50% of the embryos co-injected with gRNAa and gRNAb displayed reduced *hnf1b* expression in the pronephros in comparison to control uninjected embryos, with a strong reduction for half of them. We obtained embryos with a decreased *hnf1b* expression in response to gRNAa or gRNAb alone but these effects were not statistically significant (Fig. 6c). When *Lhx1* pronephric expression was analyzed, we did not see any effect of the gRNAs alone or in combination, highlighting the specific regulatory effect of CNS1 on *hnf1b* expression (Fig. 6d). These results indicate that CNS1 is required *in vivo* for the proper *hnf1b* pronephric expression at tailbud stage.

**Figure 6.**
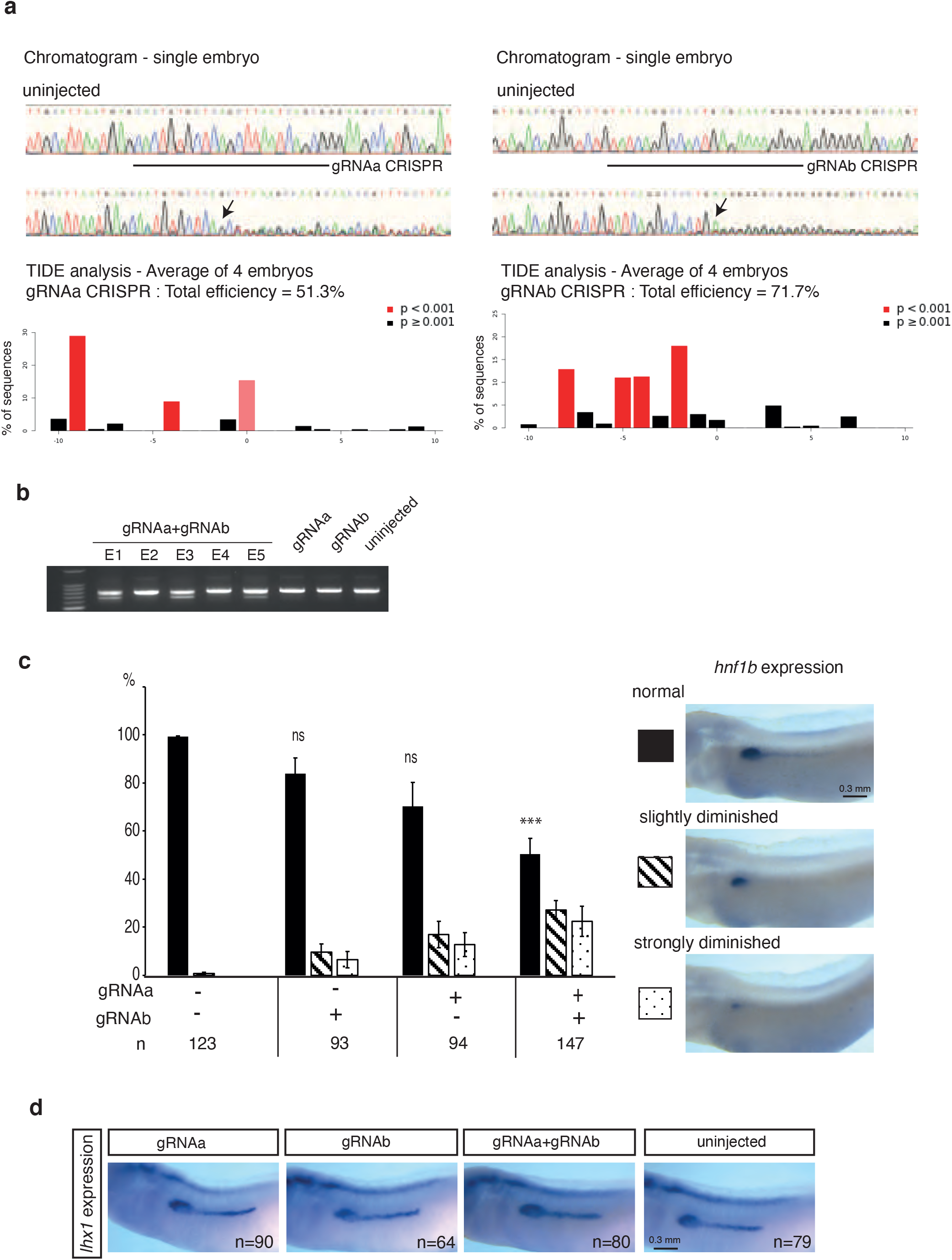
CNS1 CRISPR editing inhibits endogenous *hnf1b* pronephric expression in *Xenopus tropicalis* embryos. **(a)** GRNAa and gRNAb efficiently edit *Xenopus tropicalis* embryo DNA in two different locations in CNS1. Embryos were injected at the 1-cell stage with Cas9 protein and gRNAa or gRNAb and cultured until neurula stage. The target sequences of gRNAa and gRNAb are depicted in figure 1 (see also methods). Top panels: chromatogram showing CRISPR editing by gRNAa and gRNAb in single embryos. Lower panels: TIDE analysis of sequence trace degradation at the expected Cas9 site of DNA cleavage. Percentage of CNS1 sequence containing insertions and deletions (Indels) represented as the mean from four embryos. The use of gRNAa and gRNAb resulted in 51.3 % and 71.7 % editing efficiency, respectively **(b)** The use of gRNAa and gRNAb together efficiently leads to nucleotide sequence deletion into CNS1. Embryos were injected at the 1-cell stage with Cas9 protein, together with gRNAa, gRNAb or both. At tailbud stage, their DNA were analysed by PCR using primers allowing amplification of a DNA fragment containing the entire CNS1. Electrophoresis of amplified fragments from single embryos. A band corresponding to the wild type sequence is observed in all embryos. An additional lower band is present in 3 out of 5 embryos co-injected with gRNAa and gRNAb, indicating that a deletion occurred **(c)** *Hnf1b* pronephric expression in gRNAa, gRNAb and gRNAa-gRNAb CRISPR embryos analyzed by *in situ* hybridization at tailbud stage 28. Embryos were classified into three groups according to their *hnf1b* pronephric expression (normal, slightly diminished, strongly diminished). The histogram shows the percentage of embryos for each group. Average values from four independent experiments. Statistical significance was determined using Fisher’s exact test (comparison to the uninjected control embryos). *\*\*\*p<0.001*. n indicates the total number of analysed embryos **(d)** *Lhx1* pronephric expression in gRNAa, gRNAb and gRNAa-gRNAb CRISPR embryos analyzed by *in situ* hybridization at tailbud stage 28. No significant effect on *lhx1* expression is observed. n indicates the total number of embryos analysed from three independent experiments.

## Discussion

Enhancers confer tissue-specific gene expression required for proper development and homeostasis. In this study, we have identified an evolutionary conserved sequence (CNS1) located several kb upstream of the *Hnf1b* transcription start site (TSS) showing enhancer activity in renal tissue. We further demonstrated that CNS1 binds and is responsive to the TF Pax8, and is critical *in vivo* for *hnf1b* expression in the *Xenopus* developing pronephros.

Numerous methods have been developed with success to identify putative enhancers across entire genomes, based on evolutionary DNA sequence conservation, chromatin accessibility, histone modifications and chromatin interactions ^46–48^. Whatever the method, finding out which putative enhancers are functional, which patterns of expression they control, and how they encode these expression patterns is always a challenge. Our work highlights and confirms the power of the *Xenopus* model for comparative genomics-based approach coupled with transgenesis assay ^29^. Although *hnf1b* is expressed in other embryonic tissues such as the neural tissue and the endoderm, CNS1 drives expression in *hnf1b* pronephric territory only, indicating it is a tissue-specific enhancer. During neurulation, we did not observe any GFP reporter expression in the kidney field suggesting that CNS1 is dispensable for the initiation of *hnf1b* expression in the developing kidney but is rather critical for its maintenance. In order to have a more comprehensive understanding of the regulatory sequences governing *hnf1b* spatiotemporal expression, other approaches not depending to the strict sequence conservation should be used. It is indeed acknowledged that although genes involved in vertebrate development are unusually enriched for highly conserved non-coding sequence elements, enhancer function can be conserved without discernible sequence conservation ^49^. The number, location, and type of TF binding sites can change within an enhancer over evolution without loss of functionality ^50^. Notably, we could not find in the zebrafish genome any sequence exhibiting high similarity with CNS1. It may be because the *Xenopus* genome is much closer to that of mammalian genomes than the zebrafish genome, where many duplications and rearrangements followed by subfunctionalization of cis regulatory elements occurred after they had diverged from other vertebrates ^51,52^.

Connecting enhancers with their target genes is a major challenge. In contrast to promoters that reside in the first 1–2 kb upstream of the TSS of a gene, enhancers can be found much far away from the genes they influence. In *Xenopus tropicalis*, CNS1 is located 16.6 kb upstream of the *hnf1b* TSS and 23.6kb downstream the end of the *hnf1b* flanking gene *heatr6* (HEAT repeat containing 6 protein). Synteny is conserved in mouse and CNS1 is located 33.4 kb upstream the *Mus musculus Hnf1b* TSS, and 33.4 kb downstream the end of the *Heatr6* gene raising the question of potential role of CNS1 in *Heatr6* regulation*. Heatr6* is known to be significantly expressed in the ectoderm and neurectoderm in *Xenopus* embryos, but neither in the developing pronephros (Xenbase, https://www.xenbase.org), nor in the developing mouse kidney (GUDMAP, https://www.gudmap.org). *Heatr6* is therefore unlikely to be under the influence of CNS1. In contrast, our data clearly argue in favor of *hnf1b* being a target gene regulated by CNS1: 1/ CNS1 drives reporter expression in a subset of *hnf1b* expression territories 2/ CNS1 is responsive to Pax8, which is known to control *hnf1b* expression in the *Xenopus pronephros* ^25^ 3/ CRISPR/cas9 which allowed CNS1 editing within its native genomic context leads to a decrease of endogenous *hnf1b* expression. This last observation is striking since several lines of evidence suggest that there is ample enhancer redundancy among developmentally regulated genes. Enhancers regulating the same gene often display overlapping spatiotemporal activity, a mechanism that contribute to fine-tune both the levels and patterns of gene expression but also safeguard against genetic and environmental perturbations therefore conferring robustness to developmental processes ^53–55^. Beside pronephros expression, we found that CNS1 can promote expression of a reporter gene in domains devoid of endogenous *hnf1b* expression, such as somites (Fig. S3). This may indicate the existence of silencer elements that downregulate *hnf1b* expression in these territories. Alternatively, this ectopic expression could be a consequence of the disruption of the CNS1 genomic architecture in our assay since it is known that the genomic context plays an important role for the proper function of enhancer elements.

Enhancers bind multiple TFs to activate specific gene transcription. In the present study, we addressed the role of the AATTCAGGCAATTATCTGCAT sequence identified as a Pax8 binding site present in CNS1. Our observations allowed us to conclude that this binding site is essential for CNS1 to activate transcription in specific cellular contexts. In renal cell lines, mutation of the Pax8 binding site leads to a small decrease of CNS1 activity in MDCK cells but does not have any significant effect in IMCD3 cells. In the *Xenopus* embryo, the Pax8 binding site is required for CNS1 activity in the developing pronephros at tailbud stage when patterning of the anlage occurs. Later on, when the segmented tubule is formed, the Pax8 binding site is not required anymore for reporter gene expression, indicating that other TFs are also at work and can compensate for its mutation. Interestingly, we noticed that CNS1 harbors LEF/TCF (T-cell factor/lymphoid enhancer-binding factor) binding sites suggesting that the Wnt/beta-catenin pathway could be an input for CNS1 activity. In agreement with this hypothesis, Wnt signaling components are expressed at several stages of pronephros development (including Wnt9a, Frizzle8 and Frizzled 7 ^56^) canonical Wnt signaling activity is detected in the pronephros of the tadpole in a *Xenopus* Wnt-responsive transgenic line ^57^ and Wnt/beta-catenin signaling is required for pronephros development ^58^.

Because of their high degree of homology, both within the paired domain and in their COOH-terminal transactivation domain, Pax-2 and Pax-8 proteins share similar DNA binding motifs ^59^. Indeed, we have found that exogenous pax2 is able to activate CNS1 transcriptional activity as pax8 does, while this activation is lost when mutations are introduced in the AATTCAGGCAATTATCTGCAT consensus sequence. However, our results are not in favor of a predominant role of pax2 in CNS1 pronephric activity in the *Xenopus* embryo. Pax2 knockdown neither affects reporter gene expression driven by CNS1 nor *hnf1b* endogenous expression in the pronephros at tailbud stage. Nevertheless, we cannot rule out the possibility that pax2 may contribute to pronephric CNS1 activity *in vivo;* but that its absence may be compensated by pax8. Pax2 and pax8 functional redundancy is well documented in different tissues including kidney ^60^. During mouse kidney development, both genes act in a cooperative and dosage dependent manner as revealed by the correlation between the number of Pax2/8 alleles and the severity of renal defects ^61–63^.

Our previous work has highlighted a role for pax8 in *hnf1* expression at the early steps of pronephros development ^25^. To the best of our knowledge, among the studies aiming at the identification of Pax8 and Pax2 target genes in different tissues, *hnf1b* was not identified as direct target gene. With this work we demonstrate for the first time that *hnf1b* is directly regulated by pax8 in the developing pronephros. Whether this regulation is functionally conserved during kidney development in other vertebrates remains to be proved. In zebrafish, pax2a/8-deficient embryos fail to activate expression of *hnf1ba* in the intermediate mesoderm ^28^. In mouse, *Hnf1b* is co-expressed with *Pax2* or *Pax8* in several renal compartments of the developing kidney. Hnf1b has been shown to cooperate with Pax2 to control common pathways governing both kidney morphogenesis and ureter differentiation ^64^ but a control of *Hnf1b* expression by either Pax8 or Pax2 in the developing mouse kidney has not yet been documented. *Hnf1b* expression appears normal in *Pax2-*deficient embryos ^64^ but since Pax8 is expressed in these embryos ^60^, a definitive conclusion on *Hnf1b* regulation by Pax genes cannot be drawn. The finding that in mouse embryonic kidney but not liver or lung, CNS1 is located in open chromatin and associated with active enhancer histone marks strongly supports the hypothesis of a functional role for CNS1 in the *Hnf1b* transcriptional regulation in the developing mouse kidney. Pax2 and Pax8 have been shown to recruit activating complexes that imprint distinct patterns of histone methylation resulting in gene activation ^65,66^. Pax2 and/or Pax8 could therefore impart epigenetic changes at the *Hnf1b* locus leading to *Hnf1b* transcriptional regulation.

Variation in gene expression is a significant determinant of human disease and enhancer-related dysregulation of gene expression has been recognized as one of the main drivers in the pathogenesis of many diseases ^67^. There are several described Mendelian disorders for which specific mutations in enhancers disrupt enhancer–gene regulation and are unequivocally causal ^68^. To date, all described *HNF1B* mutations in humans are located in the HNF1B coding region or splice sites. The identification of *HNF1b* regulatory sequences and their binding TFs could contribute to the understanding of *HNF1B-*associated diseases and may open important perspectives into the diagnosis for understanding congenital anomalies of kidney and urinary tract in human.

## Methods

### Identification of conserved non-coding regions (CNSs)

We used the dynamic graphical interface ECR Browser (http://ecrbrowser.dcode.org/), a tool for visualizing and accessing data from comparisons of multiple vertebrate genomes ^69^. With minimum similarity of 70% and 100-base pair length as set parameters, sequences were filtered by conservation analysis across different vertebrate species (*Mus musculus, Homo sapiens, Rattus norvegicus, Monodelphis domestica, Gallus gallus, Xenopus tropicalis, Danio rerio*) using a mouse 157 kb genomic sequence encompassing *Hnf1* gene as a reference genome. R-Vista 2.0 (https://rvista.dcode.org) was used to identify conserved putative TFs binding sites in CNS1 ^34^. P300 and histones ChIP-seq, DNase-seq and ATAC-seq peaks were displayed using the ENCODE databases (https://www.encodeproject.org) and database from the Veenstra’s laboratory (http://veenstra.science.ru.nl/trackhubx.htm, ^30^ for *Xenopus*. General analyses and visualization were done using UCSC genome browser (https://genome.ucsc.edu/) and Ensembl genome browser (http://www.ensembl.org/).

### Electrophoretic Mobility Shift Assays

Electrophoretic mobility shift assay (EMSA) analysis was performed using the Light Shift® Chemiluminescent EMSA Kit (Thermo Scientific). Briefly, nuclear extracts were prepared from MDCK cells, while DNA-probes (Pax8-BS, Control TPO, Pax8-BS mut) were biotin-labelled at 5’End (Eurofins). Twenty μG nuclear extracts were incubated with the different biotin-labeled probes on ice for 20 minutes in a 20 μl mixture. For competition EMSA, a 10- or 100-fold molar excess of the cold probes was added to the EMSA binding reaction. For supershift assays, 1 μg of anti-Pax8 antibody (Abcam ab53490), was incubated with the nuclear extracts for additional 10 min at 4°C before adding the probe. The reaction mixture was run on 6% polyacrylamide gel in 0.5x Tris-Borate-EDTA buffer (pH 8.5) at 300 V. The gel was electro-blotted to positively charged nylon membrane using a UV-light cross-linker instrument equipped with 254 nm bulbs at 120 mJ/cm2. Biotin-labeled DNA was detected using streptavidin-horseradish conjugate and the chemiluminescent substrate contained in LightShift Chemiluminescent EMSA Kit.

### Cell culture, transfection and luciferase reporter assay

HEK293, MDCK, and IMCD3 cells were cultured in Dulbecco’s modified Eagle’s medium (DMEM) supplemented with 10% fetal calf serum. The reporter constructs used for luciferase assay were generated as the following: CNS1 was amplified by PCR from *Xenopus laevis* genomic DNA using primers 5’-CCGCTCGAGCAGAGCAGACAGGGTCTGTA −3’ and 5’-CCCAAGCTTTGACCGTCAGTTTCATGACT-3’ and inserted into pGEM®-T Easy vectors (PROMEGA). Then, CNS1 was subcloned into the pGL4.23[luc2]miniP (PROMEGA) using XhoI and HindIII to generate CNS1-Luc. Concomitantly, we introduced several point mutations variants in the Pax8 binding site using the QuikChange® II XL Site-Directed Mutagenesis Kit (Agilent Technologies) to generate the CNS1 mut-Luc vector. Primers used are 5’-CCCCTAACATACGTTGAATTAAAAATTAGGGTCCTAACCTGCATGGCTTCCCTCTGATTAAG −3’ and 5’-CTTAATCAGAGGGAAGCCATGCAGGTTAGGACCCTAATTTTTAATTCAACGTATG TTAGGGG −3’. The CNS1 and the mutated CNS1 nucleotide sequences were confirmed by sequencing. For Dual Luciferase Reporter Assay analyses, cells were plated into 12-well plates at a density of 2.5 x 10^5^ cells/well. Transfection mixtures containing Opti-MEM® Reduced Serum Media (ThermoFisher), 500 ng of reporter vector (pGL4.23[luc2]miniP called Empty-Luc, CNS1-Luc or CNS1mut-Luc), 5 ng of pCMV-renilla luciferase normalization vector and, when appropriate, 50 pg or 100pg of the expression vector (pCS2pax2, pCS2pax8VP16, pCS2pax8ΔO, Caroll and Vize, 1999, Buisson et al., 2015) were prepared. Cells were transfected using 1.5 μL of X-tremeGENE HP DNA Transfection Reagent (Roche) according to the manufacturer’s protocol. Transfected cells were washed twice in PBS, followed by the addition of 200 μl 1x passive lysis buffer (Promega, Madison, WI, USA). All values are shown as the mean+S.E.M. All experiments were repeated at least three times. Statistical significance was determined using One-Way ANOVA followed by Tuckey’s multiple comparison test (\**P*<0.05, \*\**P*<0.005, \*\*\**P*<0.001 \*\*\*\**P*<0.0001).

### Immunofluorescence

Cells on a glass slide were fixed with 3% formaldehyde at room temperature for 15 min and permeabilized with 0.2%. Triton X-100 in PBS for 5 min. Cells were then treated with 3% bovine serum albumin in PBS and incubated overnight at 4°C with the primary mouse antibody anti-Pax8 (Abcam ab53490, diluted 1/100). After washing unbound antibodies, cells were incubated with the secondary antibodies Alexa Fluor 488 anti-mouse (1/2000) along with DAPI for 1 h. Slides were washed and mounted in Mowiol mounting medium (Sigma-Aldrich). Pictures were taken using Zeiss Apotome Wide Field Microscope.

### *Xenopus* embryos, morpholino microinjection, *in situ* hybridization

*Xenopus lævis* and *Xenopus tropicalis* were purchased from the CNRS *Xenopus* breeding Center (Rennes, France). Embryos were obtained as described ^25,70^, and were raised in Modified Barth’s solution (MBS). Stages were according to the normal table of *Xenopus lævis* ^71^ and as described for *Xenopus tropicalis* ^72^. All experimental protocols and the handling of *Xenopus* were done in accordance with the European Community Directive 2010/63/UE and with the ARRIVE guidelines. They were approved by the French National Ethics Committee for Science and Health report on “Ethical Principles for Animal Experimentation” - C2EA-05 Charles Darwin - under agreement N° 27754-2021050515059177. Microinjection of morpholinos was performed according to Buisson et al, 2015. Pax2 knockdown was obtained by injection of MoPax2.2; Pax8 knockdown was obtained by co-injection of MoPax8 and MoPax8.B ^25^. Whole mount *in situ* hybridization to detect *GFP, hnf1b* and *lhx1* mRNAs were performed as in ^25^.

### Transgenic reporter assay in *Xenopus laevis*

Transgenic *Xenopus laevis* embryos were generated using the *I-SceI* meganuclease method ^44^ or the restriction-enzyme-mediated integration (REMI) method ^45^. Constructs used for transgenesis were obtained as follows. CNS1 or mutated CNS1 sequences were retrieved from CNS1-Luc and CNS1 mut-Luc plasmids using XhoI and Hind III, and were inserted upstream of a human *β-globin* basal promoter (−37 to +12) followed by an eGFP coding sequence into the beta globin-eGFP reporter vector described in Ochi et al, 2012. Manipulated embryos were cultured until the desired stage, and normally developing embryos were subjected to wholemount *in situ* hybridization for GFP mRNA detection.

### CRISPR/Cas9 editing in *Xenopus tropicalis*

GRNAa (ACTGTGCTCAGCTTAATCAGCGG) and gRNAb (TATCAGGCCACTGAGAAAAGGGG) targeting CNS1 were designed and selected for their high predicted specificity and efficiency using CRISPOR online tool (http://crispor.tefor.net/). They were purchased from Integrated DNA Technologies, as Alt-R crRNA and tracrRNAv (IDT, Coralville, IA, USA) and dissolved in duplex buffer (IDT) at 100 μM each. cr:tracrRNA duplexes were obtained by mixing equal amount of crRNA and tracrRNA, heating at 95°C for five minutes and letting cool down to room temperature. GRNA:Cas9 RNP complex was obtained by incubating 1μL of 30 μM Cas9 protein (kindly provided by TACGENE, Paris, France) with 2μL cr:tracrRNA duplex in a final volume of 10 μL of 20 mM Hepes-NaOH, 150 mM KCl, pH 7.5 for 10 min at 28°C. *Xenopus tropicalis* one-cell stage embryos were injected with 2 nL of gRNA:Cas9 RNP complex solution and were cultured to the desired stage. For co-injection, both complexes were mixed equally. Single embryo genomic DNA was obtained by digesting for 1h at 55°C in 100 μL lysis buffer (100 mM Tris-HCL pH 7.5, 1 mM EDTA, 250 mM NaCl, 0.2% SDS, 0.1 μg/ μL Proteinase K), precipitating with 1 volume of isopropanol and resuspended in 100μL PCR-grade water. The region surrounding the gRNA binding site was amplified by PCR using 5’-CCTTACGCCTTACTGAAGAGTGC-3’ as forward primer and 5’-CAGCCAGGGAACTATGTGCAA-3’ as reverse primer and PCR products were analyzed by 1.5% agarose TAE gel electrophoresis and send for sequencing. TIDE was used to determine total efficiency and insertion and deletion frequencies in the amplified gene region ^73^.

## Acknowledgements

We thank H. Ogino for kindly providing the beta globin-eGFP reporter vector. European Union’s Horizon 2020 Research and Innovation Program supported the research presented in this manuscript, under the Marie Skłodowska Curie grant agreement No. 642937 (RENALTRACT; MSCA-ITN-2014-642937) and grants from CNRS and Sorbonne Université. Laura Goea was recipient of a PhD candidate contract from ITN RENALTRACT MSCA-ITN-2014-642937. We thank S. Authier for excellent technical assistance in the maintenance of the *Xenopus* animal facility.

## Data availability

The datasets generated during and/or analyzed during the current study are available from the corresponding author on reasonable request. Public databases and bioinformatic tools used for our studies are the following : ECR Browser (http://ecrbrowser.dcode.org/) for comparisons of multiple vertebrate genomes ; R-Vista 2.0 (https://rvista.dcode.org) to identify conserved putative TFs binding sites ; ENCODE databases (https://www.encodeproject.org) to display histones ChIP-seq, DNase-seq and ATAC-seq peaks in mammals and the database from the Veenstra’s laboratory for *Xenopus* (http://veenstra.science.ru.nl/trackhubx.htm). Accessible chromatin and ChIP-seq data are visualized with the UCSC genome browser at (https://genome.ucsc.edu/) and Ensembl genome browser (http://www.ensembl.org/). Sequence alignment was performed with the short sequence aligner software ClustalW (http://www.ebi.ac.uk/Tools/msa/clustalw2/). Gene expression data are available on (https://sckidney.flatironinstitute.org), GUDMAP (https://www.gudmap.org) or Xenbase (https://www.xenbase.org).

## Supplementary Tables

**Table S1**

*In silico* identification of conserved TFs binding sites conserved in *Mus musculus* and *Xenopus tropicalis* CNS1 using r-vista 2.0 (https://rvista.dcode.org).

**Table S2**

*In silico* identification of conserved TFs binding sites conserved in *Homo sapiens* and *Xenopus tropicalis* CNS1 using r-vista vista 2.0 (https://rvista.dcode.org). In grey, TFs binding sites conserved between *Homo sapiens* and *Xenopus tropicalis* but not present in *Mus musculus*.

## Supplementary figures

**Figure S1**

Chromatin signature of a genomic region including CNS1 and CNS2 in *Homo sapiens*, *Mus musculus* and *Xenopus tropicalis*. The blue box indicates CNS1 and the orange one CNS2. **(a)** Representative DNase-seq tracks for a 123 kb genomic region in human (chr17: 37,714,024-37,837,224 – GRCh38/hg38 assembly) from ENCODE database ( https://www.encodeproject.org) for the following biological materials: embryonic kidney tissue (female embryo 108 days, ENCSR941DTJ) ; proximal tubule primary cells (ENCSR000EPW) ; adult kidney tubule primary cells (female adult 80 years, ENCSR17IWT) ; kidney epithelial primary cells (ENCSR00EOL) ; the colon carcinoma cell-line Caco-2 cell line (ENCSR000EMI) ; the epithelial lung carcinoma cell-line A549 cell line (ENCSR000ELW) ; embryo spinal cord (male embryo 96 days, ENCSR788SOI) ; neural progenitors (female embryo 5 days neural progenitor *in vitro* differentiated cells originated from H9 (ENCSR963ALV). A bottom track shows sequence conservation among the indicated vertebrate species. The observed region partly overlaps with the location of HNF1b transcript as indicated in the last bottom track. **(b)** Representative ATAC-seq and histone modifications tracks as indicated for a 65 kb genomic region in mouse (chr11: 83,810,000-83,875,000 - GRCm38/mm10 assembly) from ENCODE database (https://www.encodeproject.org) in kidney, lung and liver of E15.5 mouse embryo. **(c)** Representative ChiP-seq tracks for the indicated proteins and embryo stages for a 47 kb genomic region in *Xenopus tropicalis* (chr02: 48,882,893-48,930,436 – xT9_0 Assembly) (http://www.veenstralab.nl/trackhubx.htm) ^30^. All tracks were visualized using UCSC browser and vertical viewing range setting.

**Figure S2**

*In situ* hybridization for *pax2, pax8* and *hnf1b* in *Xenopus laevis* embryos at the indicated stages.

**Figure S3**

CNS1 activity in *Xenopus laevis* transgenic embryos generated by the *I-SceI* meganuclease method. Histograms indicating the percentage of F0 transgenic embryos generated with CNS1 -eGFP, CNS1 mut -eGFP or the control vector β-globin basal promoter, showing *GFP* expression in the indicated tissue and/or organ at stage 28 and stage 35. Results corresponding to three independent experiments. On the right, examples of *GFP in situ* hybridization performed on CNS1-eGFP transgenic embryos showing expression in the different tissue and organs. b.a.: branchial arches, p: pronephros

## Author contributions

M.U. and J-F.R. conceived and designed the study. M.U. supervised the studies and wrote the paper with S.C. Most of the experiments were performed by L.G and I.B. except CRISPR/cas9 experiments which were performed by A.E., morpholino experiments performed by V.B. and REMI transgenesis by A.C. Results were analyzed by L.G, I.B, M.P-F and R.L.B.

