## Supplementary tables and figures for "*Hnf1b* renal expression directed by a distal enhancer responsive to Pax8"

**Figure S1**

**a** -DNase seq -Human cells - ENCODE project

1 : kidney tissue (female embryo 108 days, ENCSR941DTJ)
2 : Epithelial of proximal tubule primary cells (ENCSR000EPW)
3 : Kidney tubule primary cells (female adult 80 years,  ENCSR17IWT)
4 : Kidney epithelial primary cells (ENCSR00EOL)
5 : Caco-2 cell line (ENCSR000EMI)
6 : A549 cell line (ENCSR000ELW)
7 : Spinal cord (male embryo 96 days, ENCSR788SOI)
8 : Neural progenitors (female embryo 5 days neural progenitor in vito differentiated cells originated from H9, ENCSR963ALV)

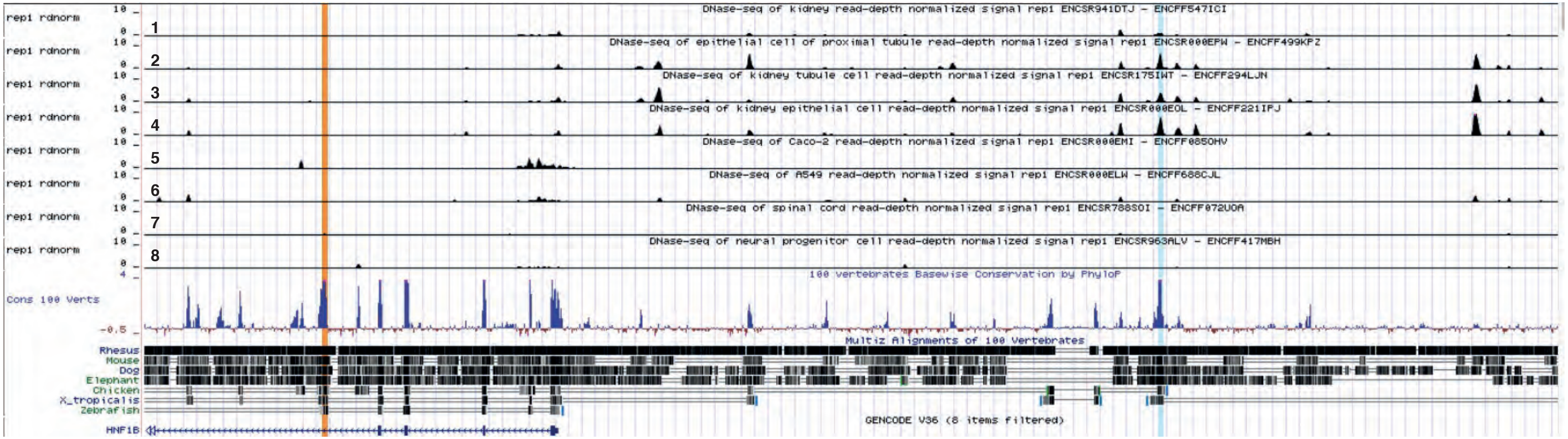


**CNS2 in orange** **CNS1 in blue**

**Figure S1**

**b** Histone modifications - Mouse embryo E15.5 - (after Gorkin D.U. *et al*., 2020)

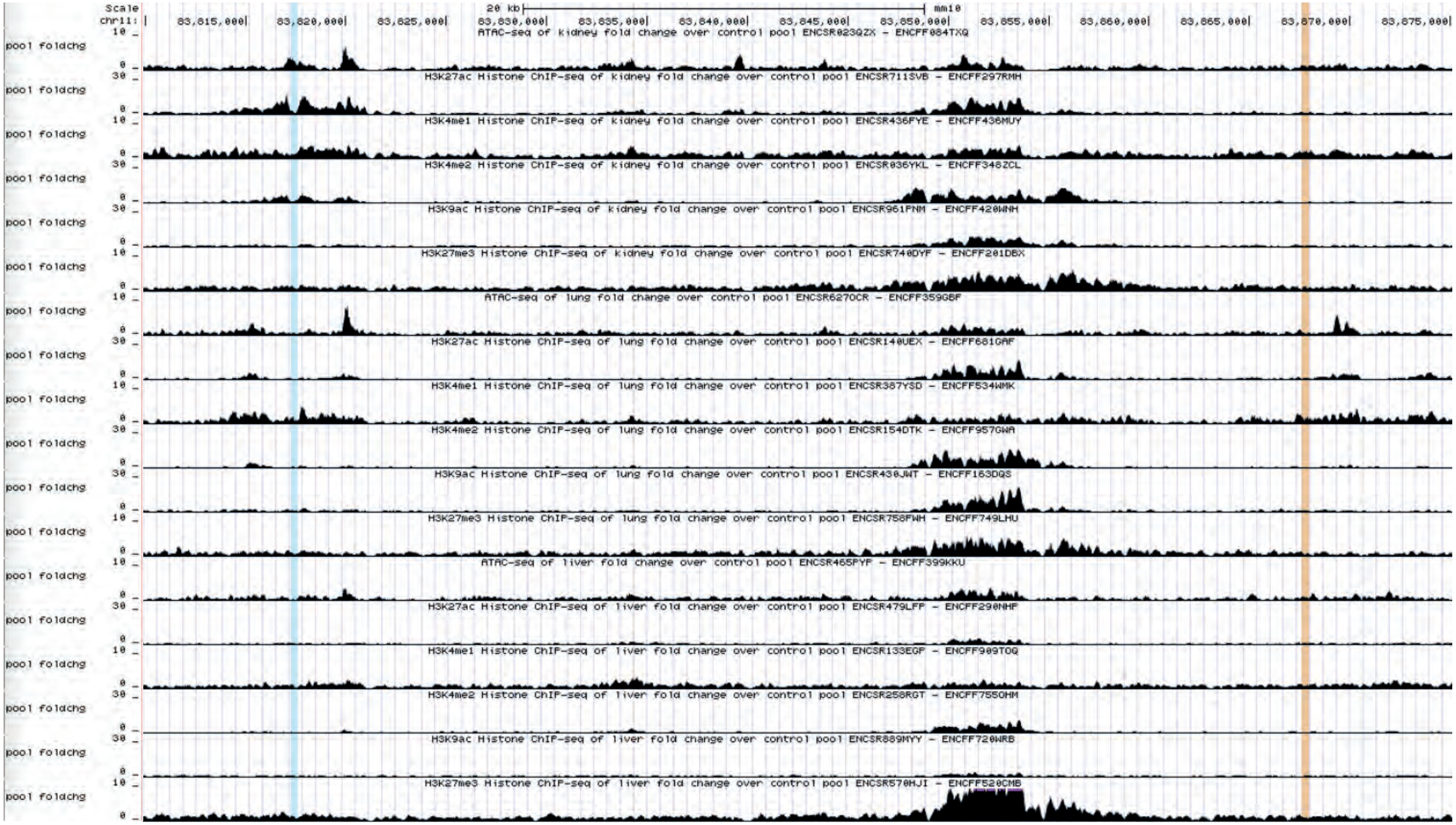


**CNS1 in blue** **CNS2 in orange**

**Figure S1**

**c** Histone modifications - Xenopus tropicalis embryos at10.5, 16 and 30 (after Hontelez S. *et al*, 2015)

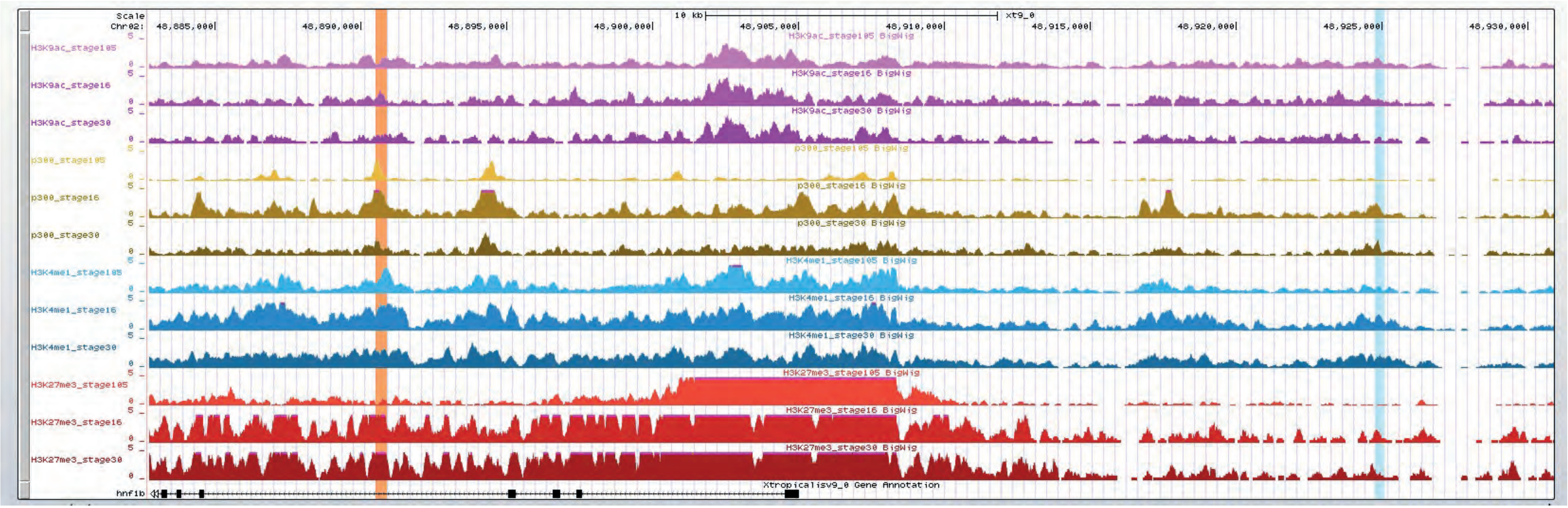


**CNS2 in orange** **CNS1 in blue**

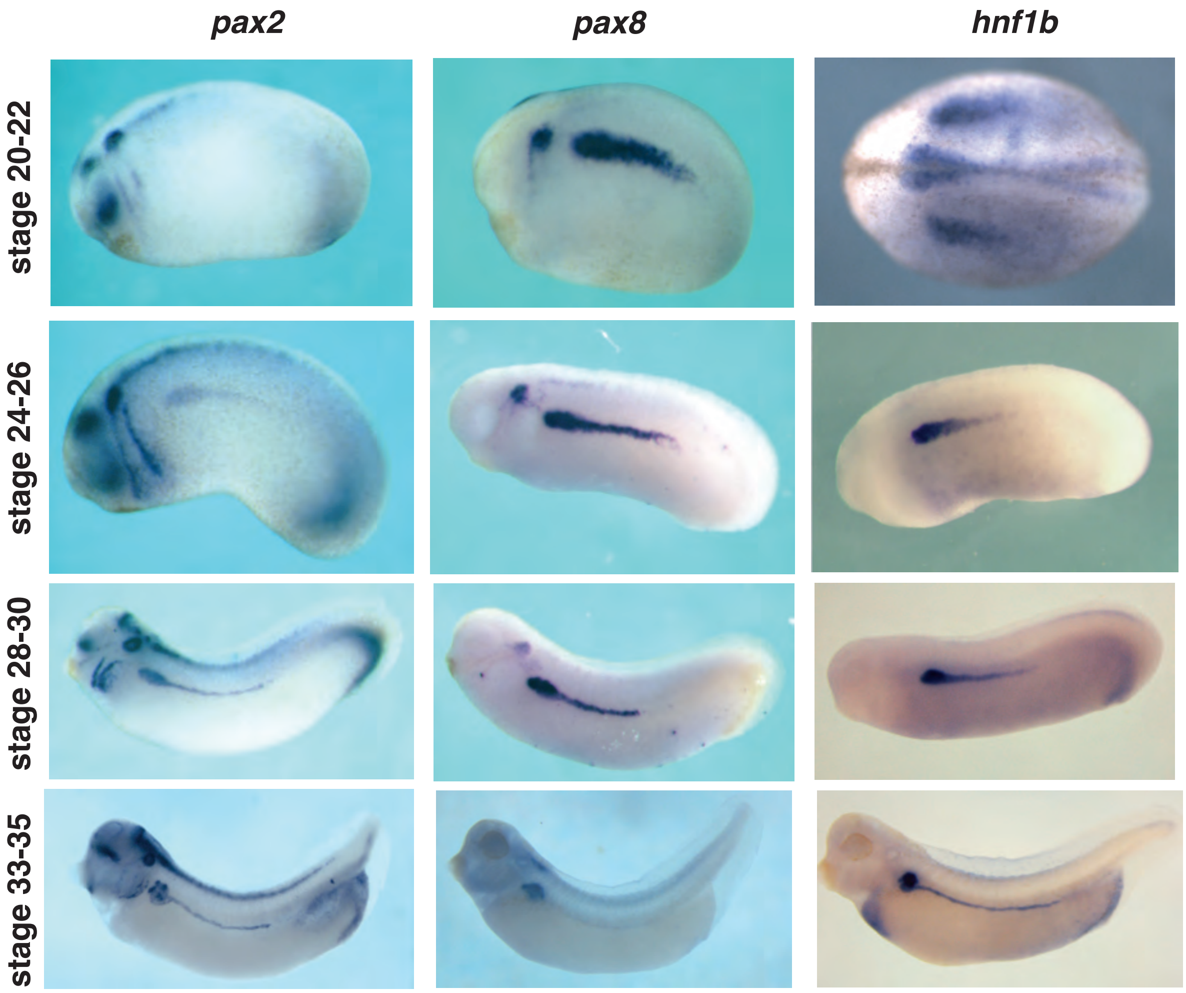


**Figure S2**

**a**

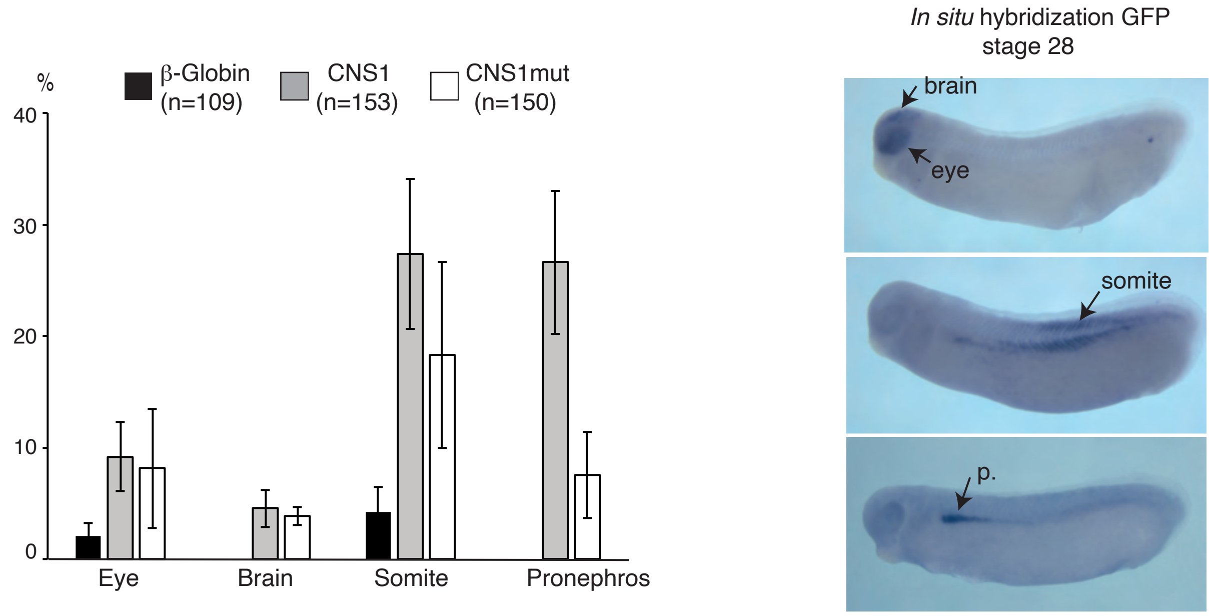

%
β-Globin (n=109)
CNS1 (n=153)
CNS1mut (n=150)
40
30
20
10
0
Eye
Brain
Somite
Pronephros
*In situ* hybridization GFP
stage 28
brain
eye
somite
p.


**b**

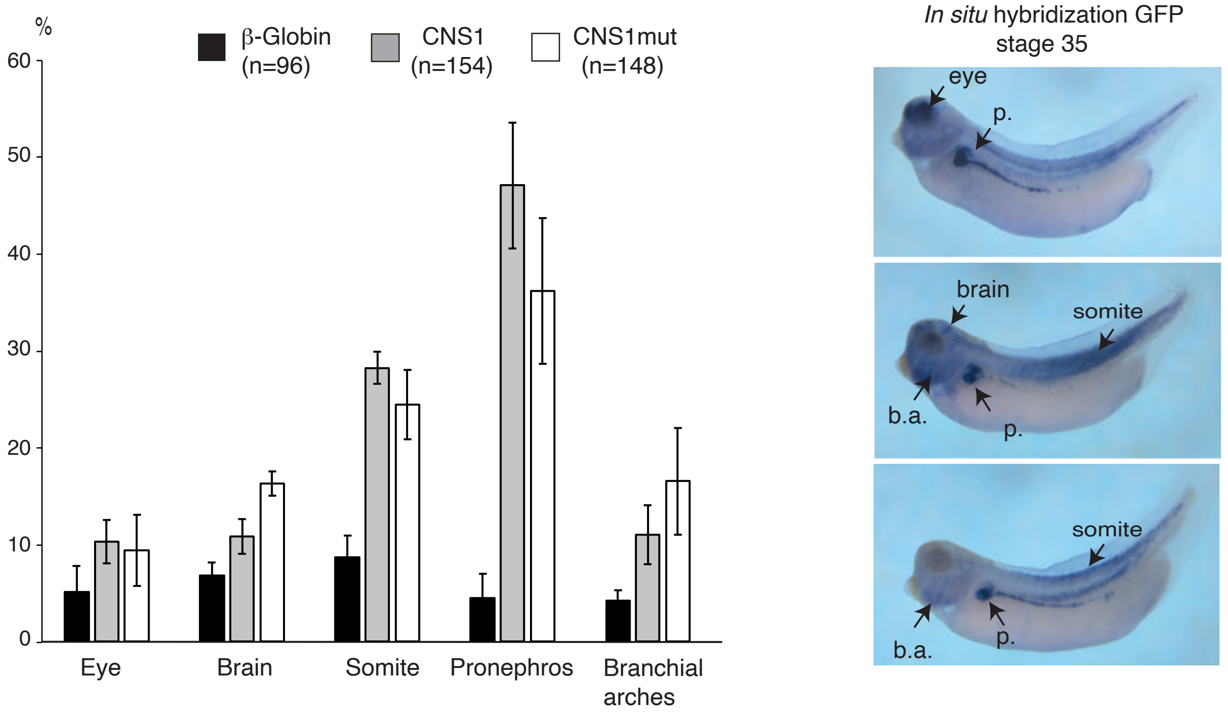

%
β-Globin (n=96)
CNS1 (n=154)
CNS1mut (n=148)
60
50
40
30
20
10
0
Eye
Brain
Somite
Pronephros
Branchial arches
*In situ* hybridization GFP
stage 35
eye
p.
brain
somite
b.a.
p.
somite
b.a.
p.


**Figure S3**

Supplementary Table 1: *In silico* identification of conserved TFs binding sites conserved in *Mus musculus* and *Xenopus tropicalis* CNS1 using R-vista 2.0.

| Conserved TF binding site | strand | Location (Mus musculus) | Sequence (Mus musculus) | Location (Xenopus tropicalis) | Sequence (Xenopus tropicalis) | Matrix score |
|---|---|---|---|---|---|---|
| V$RBPJK_01 | + | 23-33 | aaTGTGTGAAa | 24-34 | aaTGTGTGAAa | 95.00 |
| V$HMGIY_Q | + | 27-41 | tgtgaAATTTctttt | 28-42 | tgtgaAATTTcctta | 95.00 |
| V$TEF1_Q6 | - | 84-89 | CATTCC | 84-89 | CATTCC | 85.00 |
| V$ELK1_02 | + | 154-167 | tcagaAGGAAgcca | 154-167 | tcagcGGGAAgcca | 100.00 |
| V$AFP1_Q6 | | 167-177 | ATGCAGATAAT | 167-177 | ATGCAGATAAT | |
| V$PAX8_B | + | 171-188 | agataattGCCTGaattt | 171-188 | agataattGCCTGaattt | 100.00 |
| V$NKX25_02 | + | 172-179 | GATAATTg | 172-179 | GATAATTg | 100.00 |
| V$RORA1_01 | + | 211-223 | gaatcaAAGTCAc | 210-222 | gaatcaAAGTCAc | 100.00 |
| V$LEF1_Q2_ | + | 212-221 | aatCAAAGtc | 211-220 | aatCAAAGtc | 100.00 |
| V$LEF1B_01 | - | 213-219 | aTCAAAG | 212-218 | aTCAAAG | 100.00 |
| V$TCF4_Q5 | - | 213-220 | aTCAAAGt | 212-219 | aTCAAAGt | 100.00 |
| V$LEF1_Q2 | + | 214-219 | TCAAAG | 213-218 | TCAAAG | 100.00 |
| V$CMYB_01 | + | 220-237 | tcacacTGCAGTTGCtga | 219-236 | tcacacTGCAGTTGTtga | 100.00 |
| V$MYB_Q5_ | - | 225-233 | ctgcaGTTg | 224-232 | ctgcaGTTg | 100.00 |
| V$VMYB_01 | - | 226-235 | tgCAGTTgct | 225-234 | tgCAGTTgtt | 100.00 |
| V$PAX6_Q2 | - | 247-260 | tGGTTCCAGCCCAT | 246-259 | cAGTTCCAGATCAT | 85.00 |
| V$NANOG_0 | + | 253-264 | caGCCCATTGAC | 252-263 | caGATCATTGAC | 90.00 |
| V$HNF6_Q6 | - | 257-268 | ccATTGACTTtt | 256-267 | tcATTGACTTtg | 95.00 |
| V$CEBP_C | + | 261-278 | tGACTTTTGCAATTTTct | 260-277 | tGACTTTGGCAATTTTc | 95.00 |
| V$CEBPA_01 | - | 262-275 | gacttttgcAAttt | 261-274 | gacTTTGGCAAt | 95.00 |
| V$CEBP_Q2_ | - | 262-273 | gacTTTTGCAAt | 261-272 | gacTTTGGCAAt | 95.00 |
| V$CEBP_Q3 | + | 262-273 | gacttttGCAAt | 261-272 | gactttgGCAAt | 95.00 |
| V$HMGIY_Q | + | 266-280 | tttgcAATTTtcttt | 265-279 | ttggcAATTTtcttt | 95.00 |

Supplementary Table 2: *In silico* identification of conserved TFs binding sites conserved in *Homo sapiens* and *Xenopus tropicalis* CNS1 using R-vista 2.0. In grey, TFs binding sites conserved between *Homo sapiens* and *Xenopus tropicalis* but not present in *Mus musculus*

| Conserved TF binding site | strand | Location (Homo sapiens) | Sequence (Homo sapiens) | Location (Xenopus tropicalis) | Sequence (Xenopus tropicalis) | Matrix score |
|---|---|---|---|---|---|---|
| V$HMGIY_Q | - | 27-41 | aaagaAAATTgcaaa | 265-279 | ttggcAATTTtcttt | 95.00 |
| V$CEBP_C | - | 29-46 | agAAAATTGCAAAAGT | 260-277 | tGACTTTGGCAATTTT | 95.00 |
| V$CEBPA_01 | + | 32-45 | aaaTTgcaaaagtc | 261-274 | gactttggcAAttt | 95.00 |
| V$CEBP_Q3 | _ | 34-45 | aTTGCaaaagtc | 261-272 | gactttgGCAAt | 95.00 |
| V$CEBP_Q2 | + | 34-45 | aTTGCAAAAgtc | 261-272 | gacTTTGGCAAt | 95.00 |
| V$HNF6_Q6 | + | 39-50 | aaAAGTCAATgg | 256-267 | tcATTGACTTtg | 95.00 |
| V$FXR_IR1_ | + | 41-53 | AAGTCAATGGCCt | 253-265 | aGATCATTGACTT | 90.00 |
| V$NANOG_0 | - | 43-54 | GTCAATGGCCtg | 252-263 | caGATCATTGAC | 90.00 |
| V$GCNF_01 | - | 46-63 | aaTGGCCTGGAACTA | 243-260 | ttCCAGTTCCAGATCAt | 90.00 |
| V$PAX6_Q2 | + | 47-60 | ATGGCCTGGAACTa | 246-259 | cAGTTCCAGATCAT | 90.00 |
| V$CMYB_01 | - | 70-87 | tcaGCAACTGCAgtgtg | 219-236 | tcacacTGCAGTTGTtg | 100.00 |
| V$VMYB_01 | + | 72-81 | agcAACTGca | 225-234 | tgCAGTTgtt | 100.00 |
| V$MYB_Q5_ | + | 74-82 | cAACtgcag | 224-232 | ctgcaGTTg | 100.00 |
| V$RORA1_0 | - | 84-96 | gTGACTTtgattc | 210-222 | gaatcaAAGTCAc | 100.00 |
| V$LEF1_Q2_ | - | 86-95 | gaCTTTGatt | 211-220 | aatCAAAGtc | 100.00 |
| V$TCF4_Q5 | + | 87-94 | aCTTTGAt | 212-219 | aTCAAAGt | 100.00 |
| V$LEF1_Q2 | - | 88-93 | CTTTGA | 213-218 | TCAAAG | 100.00 |
| V$LEF1B_01 | + | 88-94 | CTTTGat | 212-218 | atCAAAG | 100.00 |
| V$STAT3_02 | + | 91-98 | tgaTTCCc | 208-215 | gGGAAtca | 100.00 |
| V$GATA1_0 | - | 95-107 | tcccCTATCataa | 199-211 | gtatGATAGggga | 100.00 |
| V$GATA3_0 | - | 97-105 | ccctATCat | 208-215 | gGGAAtca | 100.00 |
| V$LMO2CO | - | 97-105 | cccTATCat | 201-209 | atGATAggg | 100.00 |
| V$PAX8_B | - | 119-136 | aaattCAGGCaattatct | 171-188 | agataattGCCTGaattt | 100.00 |
| V$NKX25_0 | - | 128-135 | cAATTATC | 172-179 | GATAATTg | 100.00 |
| V$AFP1_Q6 | + | 130-140 | ATTATCTGCAT | 167-177 | ATGCAGATAAT | 100.00 |
| V$ELK1_02 | - | 140-153 | tggcTTCCTtctga | 154-167 | tcagcGGGAAgcca | 100.00 |
| V$CMAF_01 | - | 175-193 | ttaatgcaTCAGCAcaga | 114-132 | ctttgTGCTGTtgcattaa | 95.00 |
| V$TEF1_Q6 | + | 218-223 | GGAATG | 84-89 | CATTCC | 85.00 |
| V$HMGIY_Q | - | 266-280 | aaaagAAATTtcaca | 28-42 | tgtgaAATTTcctta | 95.00 |
| V$RBPJK_01 | - | 274-284 | tTTCACACAtt | 24-34 | aaTGTGTGAAa | 95.00 |
| V$TBX5_Q5 | + | 275-284 | tTCACACATt | 24-33 | aATGTGTGAa | 95.00 |
